# Genome-wide identification and expression analysis of bHLH transcription factors reveal their putative regulatory effects on petal nectar spur development in *Aquilegia*

**DOI:** 10.1101/2022.04.20.488976

**Authors:** Xueyan Li, Hui Huang, Zhi-Qiang Zhang

## Abstract

The basic helix-loop-helix (bHLH) transcription factors (TFs) control a diversity of organ morphogenesis involved in cell division and cell expansion processes. The development of petal nectar spur, which plays important roles in plant-pollinator interaction and adaptive radiation, comprised cell division and cell expansion phases in *Aquilegia*. Here, we conducted a genome-wide identification of the bHLH gene family in *Aquilegia* to determine the characteristics and the expression profiles of this gene family during the development of petal nectar spur. A total of 120 *AqbHLH* proteins were identified from the *Aquilegia coerulea* genome. The phylogenetic tree showed that *AqbHLH* members were divided into 15 subfamilies, among which S7 and S8 subfamilies occurred marked expansion. Nineteen residues with conservation of more than 50% were found in the four conserved regions. The publicly RNA-Seq data and qRT-PCR results showed that *AqbHLH027, AqbHLH083, AqbHLH046,* and *AqbHLH092* would be associated with the development of petal nectar spur by regulating cell division and cell cycle in phase I. While *AqbHLH036* might participate the spur cell elongation and cell expansion in phase Ⅱ. This study provides useful insights for further probing on the function of *AqbHLH* TFs in the regulation of petal nectar spur development.

## 1. Introduction

In general, the cell division decides the total number of cells in an organ, whereas cell expansion determines the size of cells (Wang et al., 2018a). The complex spatiotemporal patterns of them determine the size and shape of an organ from primordium, such as leaves, flowers, and fruits (Fox et al., 2018; Ripoll et al., 2018; John and Qi, 2008). For example, *VvCEB1* controls grape size via regulating cell expansion and *OsRac1* controls rice grain size and yield by regulating cell division (Zhang et al., 2019). Given their effects on agricultural production and agronomic characters either alone or in combination, it is vital to reveal the molecular mechanism of cell division and/or cell expansion for molecular breeding and cultivating new varieties. Transcription factors (TFs), form a complex system directs expression of the genome in universal or special space and time, can bind to specific *cis*-acting elements in the promoters regions of downstream target genes, in turn, involved in many biological processes. The basic helix-loop-helix (bHLH) TF is the second-largest gene family in plants following MYB gene family (Feller et al., 2011). The bHLH gene family incorporates two conserved regions, a helix-loop-helix region (HLH region) related to the formation of homodimers or heterodimers and a basic region determined the DNA binding specificity (Toledo-Ortiz et al., 2003). The bHLH TFs are essential in transcriptional networks systems in growth, development and metabolic process, such as flowering and fruits development (Ito et al., 2012, Li et al., 2019), plant morphological development (Arai et al., 2019) and response to abiotic stress (Kavas et al., 2016). bHLH TFs control a diversity of plant organ morphogenesis associated with cell division and cell expansion have been confirmed. For instance, *AtbHLH137/CKG* regulates cell cycle progression and organ growth (Park et al., 2021). *AtbHLH40*/*INDEHISCENT* and *AtbHLH24*/*SPATULA* interact to control the development of carpel margin tissues (Heisler et al., 2001; Girin et al., 2011). PIFs, novel Myc-related bHLH TFs, regulate cytokinesis during apical hook development in specific space and time (Zhang et al., 2021). *AtbHLH45/MUTE* terminates the asymmetric cell division and differentiation of stomata (Pillitteri et al., 2007). *AtbHLH83*/*RHD6* regulates the root hair initiation and elongation (Feng et al., 2017). According to their evolutionary relationships and conservation of specific sequences, the bHLH gene family is generally divided into 15 ∼ 31 subfamilies in which members usually have similar structures and functions (Pires and Dolan, 2010). Thus bHLH gene family is a particularly promising target for better understanding of how cell division and/or cell expansion shape some agronomic or horticultural characters.

The *Aquilegia* is an ideal fora model system for the evolution and ecology of petals, the genus also has long been noted for the diversity of shape and color of its petals. Petal nectar spur, a hollow extension of the petal that is often filled with nectar to attract pollinators, is a key innovative trait related to the re-initiation of the meristematic activity program and function to promote diversification rates in multiple angiosperm lineages (Shan et al., 2019; Hodges and Arnold, 1995). Petals spurs have a remarkable diversity of length in *Aquilegia* species (Puzey et al., 2012; Zhou et al., 2019), and the spur develops with two distinct phases corresponding to cell division (Phase I) and cell elongation (Phase II). During phase Ⅰ, it occurs diffusely throughout the petal primordium but ceases completely when the spur length is ∼7-10 mm. Subsequently, spur enters phase Ⅱ, in which lengthening of the spur is entirely dependent on anisotropic cell elongation (Puzey et al., 2012; Yant et al., 2015; Ballerini et al., 2019). Previous studies have found that *POPPVICH*, a C2H2 TF, plays a central role in controlling the presence or absence of spur by regulating cell proliferation during phase Ⅰ was confirmed in *A. coerulea* (Ballerini et al., 2020)*. ARF6* and *ARF8,* members of the ARF family of transcription factors, are responsible for the variation of spur length by regulating the cell elongation in *A. coerulea* (Zhang et al., 2020). It has been reported that the *Arabidopsis ARF8* protein represses petal growth by interacting with bHLH family transcription factor BPEp, indicating that members of bHLH family might be involved in spur developments (Varaud et al., 2011). However, the regulatory mechanism of the petal nectar spur development in *Aquilegia* is extremely complex and needs to be further explored. To date, the characterization and expression analyses of the bHLH TFs and their roles in the development of petal nectar spur in *Aquilegia* have not been reported.

In this study, we performed a comprehensive genome-wide level analysis of bHLH TFs in *Aquilegia,* including gene identification, phylogenetic analysis, characterization of conserved motifs, gene structure, chromosomal location, promoter *cis*-elements and protein-protein interactions. Moreover, we also investigated the expression of all identified *AqbHLHs* in different floral organs of *Aquilegia coerulea* and compared the expression pattern of these genes in spurred and spurless *Aquilegia* species at different developmental stages. To verify the results of RNA-seq analysis, we selected 20 *AqbHLH* genes included five potential key genes for expression analysis by using quantitative real-time PCR (qRT-PCR). This study provides useful information for further research on the function of *AqbHLH* TFs in the regulation of petal nectar spur development.

## 2. Materials and methods

### 2.1 Identification of bHLH genes and bHLH conserved domain in *A. coerulea*

The genome and protein sequences of the *A. coerulea* were downloaded from Phytozome (https://phytozome.jgi.doe.gov/) (Yant et al., 2015). The bHLH protein sequences of *A.thaliana* were retrieved from PlantTFDB (http://planttfdb.gao-lab.org). We used *A. thaliana* bHLH proteins sequences as query sequences to gain the *A. coerulea* candidate sequences by using the BLASTP program (e < 10^-5^). We also applied to the identification of homeobox (PF00010) in the Pfam 32.0 database (http://pfam.xfam.org/) with the HMMER (version 3.2.1) (http://hmmer.org/download.html). All of the *AqbHLH* proteins full-length amino acid sequences were aligned with MEGA X using default parameters. According to the characteristics of bHLH domain in other plants and the results of conserved motifs, we carried out the bHLH conserved domain in *A. coerulea*. The results were visualized by using Weblogo online software (http://weblogo.threeplusone.com/) and GeneDoc (https://github.com/karlnicholas/GeneDoc).

### 2.2 Phylogenetic analysis

To investigate phylogenetic relationships of *AqbHLHs*, multiple sequence alignment of the bHLH domains of 162 *A.thaliana* (*AtbHLHs*) and 120 *A.coerulea* (*AqbHLHs*) bHLH proteins was performed by MAFFT software using default parameters (http://mafft.cbrc.jp/alignment/server/). Then, a Neighbor-Joining (NJ) phylogenetic tree was constructed by MEGA X, with the bootstrap values of 1000 replicates. We visualized the phylogenetic tree with iTOL (https://itol.embl.de/) (Letunic and Bork, 2016).

### 2.3 Protein properties, conserved motifs, gene structures, cis-elements and PPI analyses of *AqbHLH* genes

We computed the protein properties of the *AqbHLHs*, molecular weight (MW) and isoelectric point (PI) using ExPASy-ProtParam (http://web.expasy.org/protparam/). And predicted the subcellular localization of *AqbHLHs* with ProComp 9.0 (http://linux1.softberry.com). MEME (https://meme-suite.org/meme/tools/meme) was used to analyze the conserved motifs, with 15 motifs and width range from 6 to 150 bp (Bailey et al., 2006). To analyze the exon and intron of the *AqbHLHs* genes, the Gene Structure Display Server (GSDS) (http://gsds.cbi.pku.edu.cn/) was used. The DNA sequences (2000 bp) upstream of the initiation codon for each *AqbHLH* gene was extracted, and the cis-elements were predicted with PlantCARE (http://bioinformatics.psb.ugent.be/webtools/plantcare/html/). The information of elements was listed in **Supplementary Table 1**. Visualized gene structures, cis-elements with TBtools includes (Chen et al., 2020). We constructed protein-protein interaction network (PPI) of key *AqbHLHs* by using the homology in *Arabidopsis* online (https://cn.string-db.org/) and modified it using AI.

### 2.4 Expression analysis of *AqbHLHs* in spur development using public RNA-seq data

We downloaded the raw RNA-seq data of different development (spur at the 1, 3 and 7mm stages) and different tissue (spur and blade) of *A. coerulea*, *A. ecalcarata* (a spurless species), and spurred taxa species (*A. chrysantha, A. formosa, A. sibirica*) from the NCBI Sequence Read Archive (SRA) database (https://www.ncbi.nlm.nih.gov/sra/) under the BioProject number PRJNA270946 (Yant et al., 2015) and PRJNA515749 (Ballerini et al., 2019). We mapped reads to the *A.coerulea* reference genome by using Hisat2. The Stringtie was used to analyze gene expression level including splice variants, and the Transcripts Per Kilobase of exon model per Million (TPM) value was used to normalize gene expression level (Pertea et al., 2016; Zhao et al., 2020). Differentially expressed genes (DEGs) were defined with a threshold of FoldChange ≥ 2 and FDR ≤ 0.05 by using edgeR (Robinson et al., 2010). We maintained genes from our analysis with TPM > 1 in all samples, and we performed the heat maps of gene confidence expression by using R software (version 4.0.2).

### 2.5 Plant material and qRT-PCR analysis

We cultivated *Aquilegia coerulea* at Yunnan University following to the method described in Sharma and Kramer (2013). The length of spur is 1mm, 3mm and 7mm, named 1mm stage, 3mm stage and 7mm stage, consistent with Yant et al. described (2015). S-11C is described in Min and Kramer (2013). We collected the blade tissue and spur tissue at per stage. All the samples collected at least three biological replicates. All samples were immediately frozen in liquid nitrogen and stored at -80℃ for RNA extraction.

We extracted and isolated the total RNA using the HiPure Total RNA kit (Magen, R4111, China). RNA reverse transcription was carried out with the FastKing RT Kit with gDNase (TIANGEN, KR116, China). The resulting cDNAs were diluted at 1:10 as templates. qRT-PCR was conducted using the PrefectStart Green qPCR SuperMix, Passive Reference Dye (TRANGEN, China) in the QuantStudio (TM) 7 Flex Real-Time PCR System. At least three biological replicates per sample, with three technical replicates for each biological replicate. We calculated relative gene expression values using the comparative CT (2^-△△CT^) method (Livak and Schmittgen, 2001), with the *AqIPP2* gene being used as an internal control (Sharma and Kramer, 2013). And the average expression value of blade tissue at 1mm stage was used as a secondary standardization for each gene. The primers employed for qRT-PCR were designed with the Primer6 and DNAMAN. All primers are listed in **Supplementary Table 2**.

## 3. Results

### 3.1 Identification of bHLH gene family members in *A. coerulea*

A total of 188 bHLHs were detected in *A. coerulea* by using the BLASTP program. After discarding these sequences with incomplete open reading frames and redundant splice variants, 120 bHLHs were retained and designated as *AqbHLH001-120* according to their chromosomal positions (**Supplementary Table 3**). Subsequently, the protein sequences of *AqbHLHs* have been used to analyze protein physicochemical properties. The results showed that the *AqbHLHs* varied in length from 74 (*AqbHLH040*) to 955 amino acid (aa) (*AqbHLH003*) with average 351aa, isoelectric points ranged from 4.50 (*AqbHLH107, AqbHLH112*) to 10.83 (*AqbHLH003*), and their molecular weight ranged from 8.27 (*AqbHLH040*) to 104.65 (*AqbHLH003*) kDa. Subcellular localization analysis demonstrated that over 82.5% *AqbHLHs* were located in the nucleus (**Supplementary Table 3**).

### 3.2 Domain recognition and phylogenetic analysis of *AqbHLH* proteins

To investigate the amino acid sequence features, we performed multiple sequence alignment analyses of 120 bHLH domains of candidate *AqbHLHs*. Our results showed that the length of *AqbHLH* domains ranged from 54 to 62aa (**Figure 1**). The *bHLH* domain of each putative *AqbHLH* protein showed four conserved regions, including the basic region (1-13 located), the helixⅠregion (14-28 located), the loop region (29-35 located) and the helixⅡregion (36-62 located). The *bHLH* domain was generally conserved in *A. coerulea*, with 19 residues being identified as having conservation of more than 50%, including five located in the basic region, one in the loop region, four in the helix Ⅰ region and nine in the helix Ⅱ region. However, *AqbHLH034*, *AqbHLH035*, *AqbHLH040*, *AqbHLH041*, *AqbHLH052*, *AqbHLH057*, *AqbHLH066*, *AqbHLH094*, and *AqbHLH102* have no Arginine in the basic region, and are similar with the atypical *AtbHLH* proteins **(Figure 1**). In addition, compared with the conservatism of two helixes, the loop region which consisted of 0-7aa showed variation in the number and sequence of the amino acids in most of the *AqbHLHs*.

**Figure 1.**
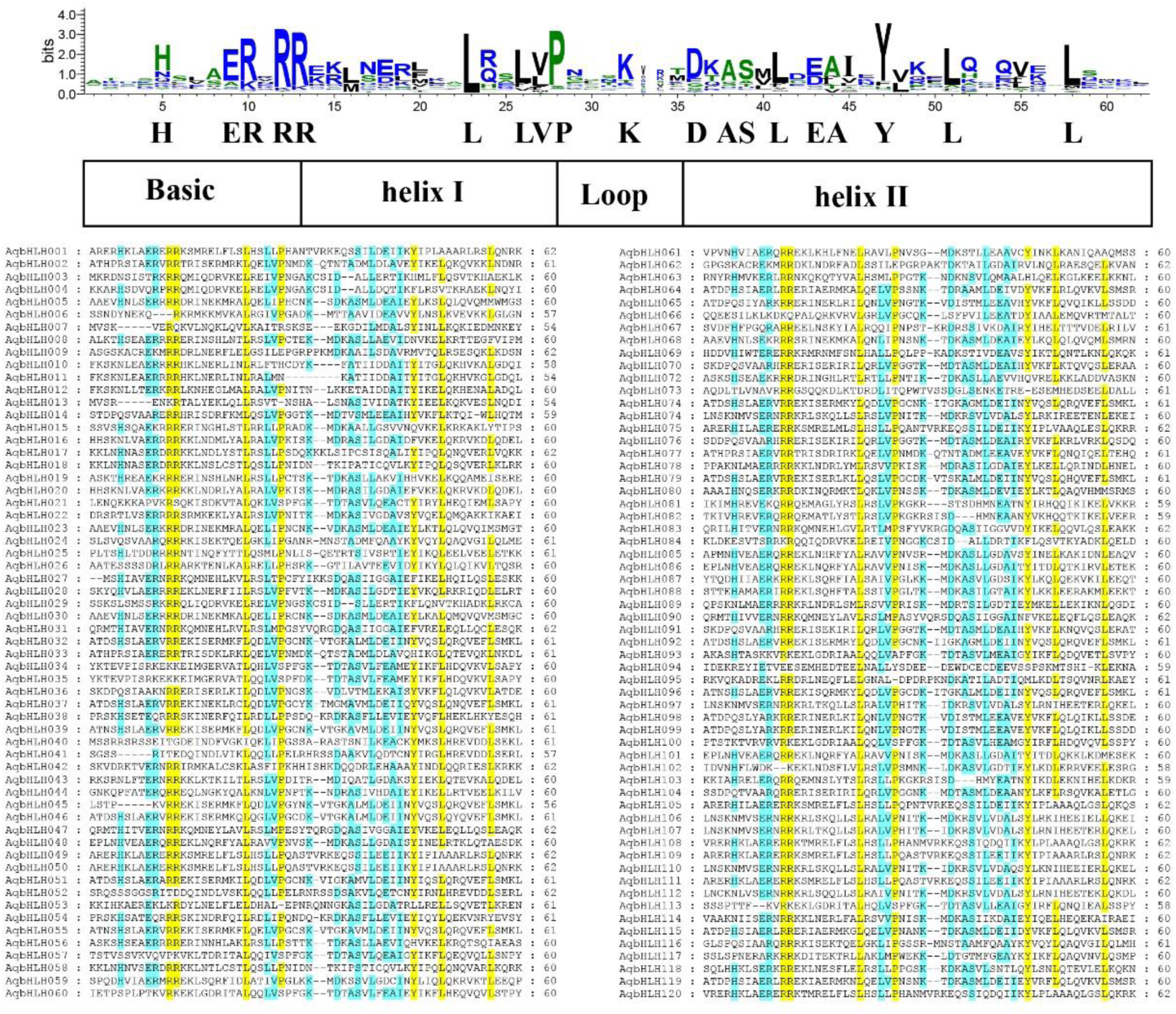
Conserved residue analysis and multiple sequence alignment of *AqbHLH* proteins. The blue background represents the residues with a consensus ratio greater than 50%, the yellow background represents the residues with a consensus ratio greater than 85%.

To classify and investigate phylogenetic relationships of the *AqbHLHs*, we constructed a Neighbour-Joining (NJ) phylogenetic tree based on the *bHLH* domains from *A. coerulea* (120) and *A. thaliana* (162). Overall, those *bHLHs* were divided into 15 subfamilies and *AqbHLH* genes were assigned to each subfamily (**Figure 2**). Notably, *AqbHLHs* were obviously expended in subfamily 7 (S7) and S8. The S7 subfamily included only one *AtbHLHs* (AT1G49770.1) and ten *AqbHLHs* genes. The S8 subfamily included ten *AtbHLHs* and 18 *AqbHLHs* members. Of them, nine *AqbHLH* genes were clustered in the subgroups with the *AtbHLH* (AT1G10610.1) in S8 subfamily, implying similar functions of them.

**Figure 2.**
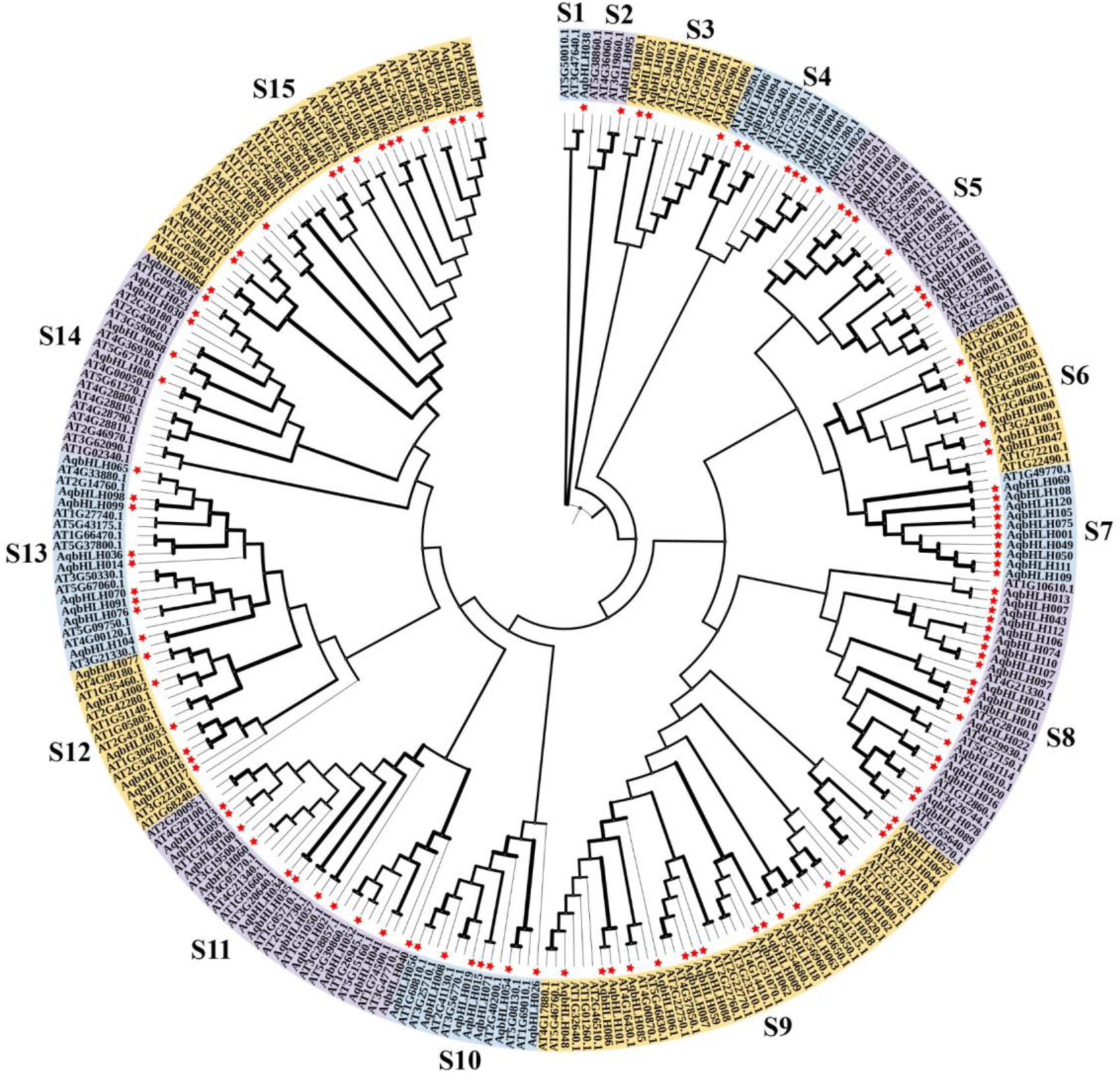
Phylogenetic tree analysis of *Aquilegia coerulea* and *Arabidopsis thaliana* bHLH proteins. The red five-pointed stars indicate the bHLH proteins from *A. coerulea*. The width of the line indicates the bootstrap value.

### 3.3 Motifs, genes structure and promoter cis-acting elements analysis

The conserved motifs and gene structures of the *AqbHLHs* were analyzed. Most members of *AqbHLHs* contained motif 1 and motif 2. However, the *AqbHLH040* and *AqbHLH066* lack motif 1 and the *AqbHLH011* lacks motif 2. Motif 3 and 4 were exclusively included in a large part of the members in S7 subfamily. The motif 5, 10 and 15 showed solely in S8 subfamily. And S5, S6, S7, S10, S14 and S15 shared motif 6. The number of exons in the 120 *AqbHLH* genes ranged from 1-13, of which *AqbHLH076* and *AqbHLH117* had only one exon. As expected, it indicated that most genes with close evolutionary relationships had similar intron-exon organizations. For example, all the members of subfamily S15 have more than six exons. Four members in subfamily S9 and S13, respectively, were intronless. These results showed that members of the same group possessed similar motifs and gene structures **(Figure 3)**.

**Figure 3.**
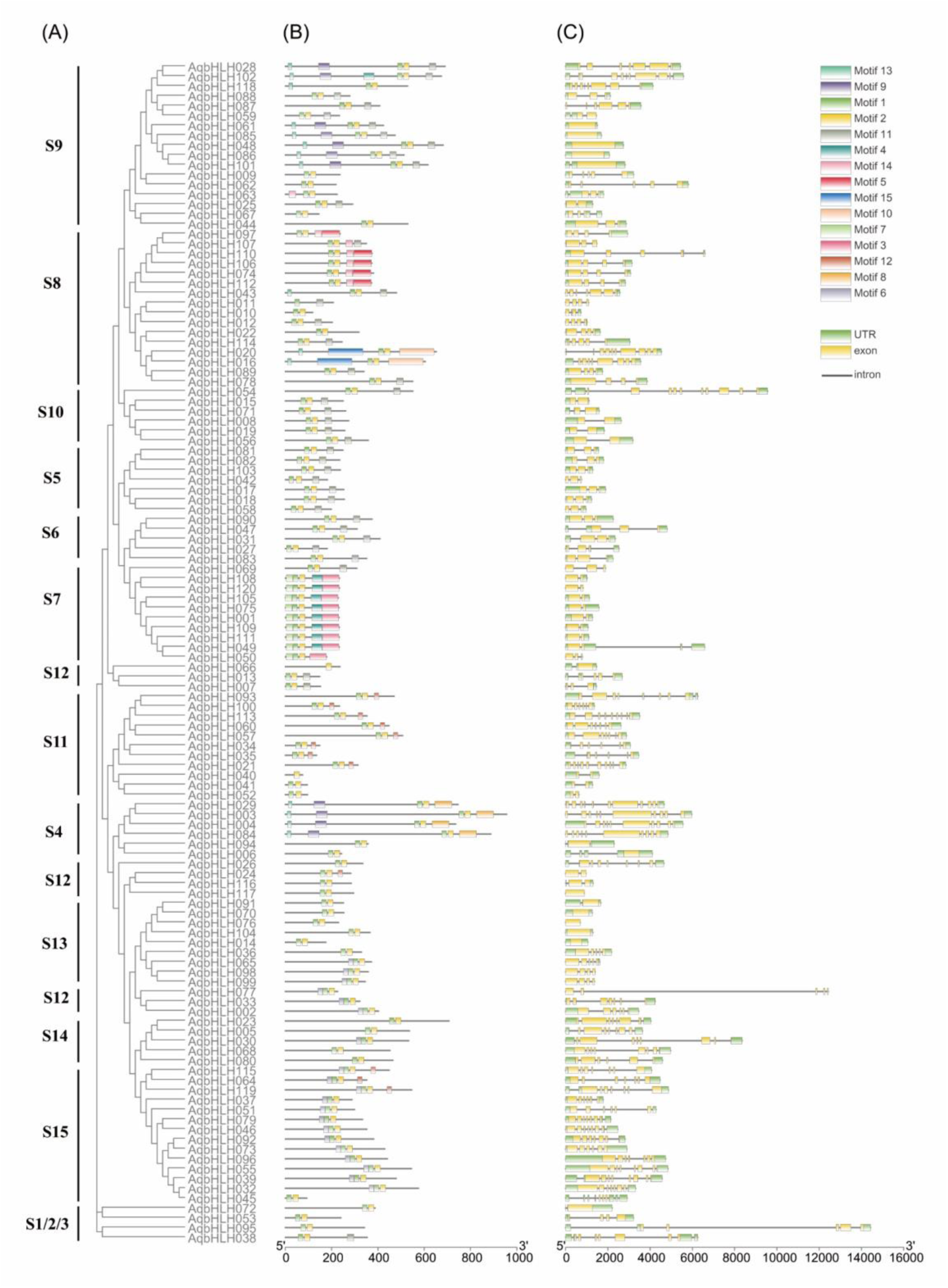
The phylogeny **(A)**, the 15conserved motifs **(B)** and gene structures **(C)** of *AqbHLH* domains, proteins and genes.

To further research the regulatory mechanism of the *A. coerulea bHLH* TFs, we extracted the upstream 2000bp sequence of 120 *AqbHLH* genes from the *A. coerulea* genome. A total of thirty type *cis*-acting elements were identified, belonging to five classes related to environment and stress, six classes related to hormone-responsive elements and five classes related to development. The light-responsive elements were the most common *cis*-acting element. Each gene had more than three light-responsive elements except for *AqbHLH066.* The *AqbHLH105* had the most light-related elements (20). The elements related to anaerobic induction were also abundant in *AqbHLH* TFs. A total of 106 *AqbHLH* genes contain anaerobic induction elements, accounting for 88% of the total number of genes. We also found that many elements were related to abscisic acid, auxin, MeJA, gibberellin, zein metabolism regulation and salicylic acid, suggesting that these *AqbHLH* genes may be involved in hormone signaling. Among the hormone-responsive elements, abscisic acid and MeJA were abundant. For instance, *AqbHLH022* and *AqbHLH100* had nine abscisic acid responsiveness elements and *AqbHLH069* had 12 MeJA responsiveness elements. *AqbHLH050* has eight auxin responsiveness elements which are more than others, suggesting that that gene may be related to auxin signaling. There were few *cis*-elements associated with development, such as meristem development, circadian control and endosperm growth. Forty-three genes had meristem expression elements and *AqbHLH 117* was the most with six. Additionally, a small part of *AqbHLH* promoters with cell cycle regulation elements, including *AqbHLH022, 029, 033, 053* and *083*. That suggested that these genes may be associated with cell cycle regulation **(Figure 4)**.

**Figure 4.**
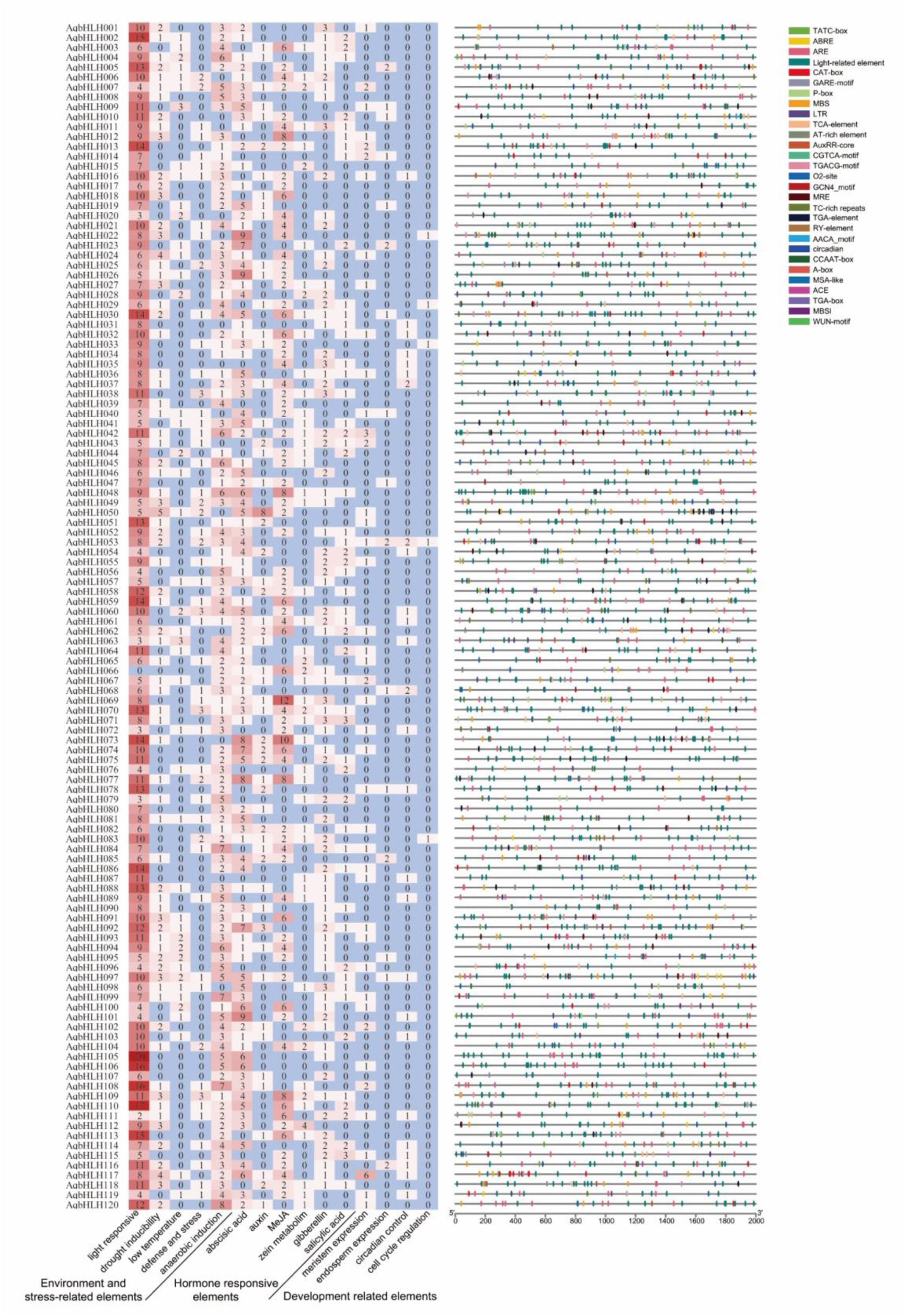
The analysis of promoter cis-acting elements for *AqbHLHs*.

### 3.4 Chromosomal distribution and synteny analysis of *AqbHLH* genes

According to the *A. coerulea* gene annotation, we localized the *AqbHLH* genes on the seven *A.coerulea* chromosomes and found that 119 *AqbHLH* genes were unevenly distributed across all of them, except for the *AqbHLH120* due to the missing annotation information. The least members were located on chromosome Ⅳ with 4 members, and chromosome Ⅴ had the most *AqbHLH* genes with 24 members. The distribution of *AqbHLH* genes is relatively tight near the ends of each chromosome, and near the centromere is sparse (**Figure 5**).

**Figure 5.**
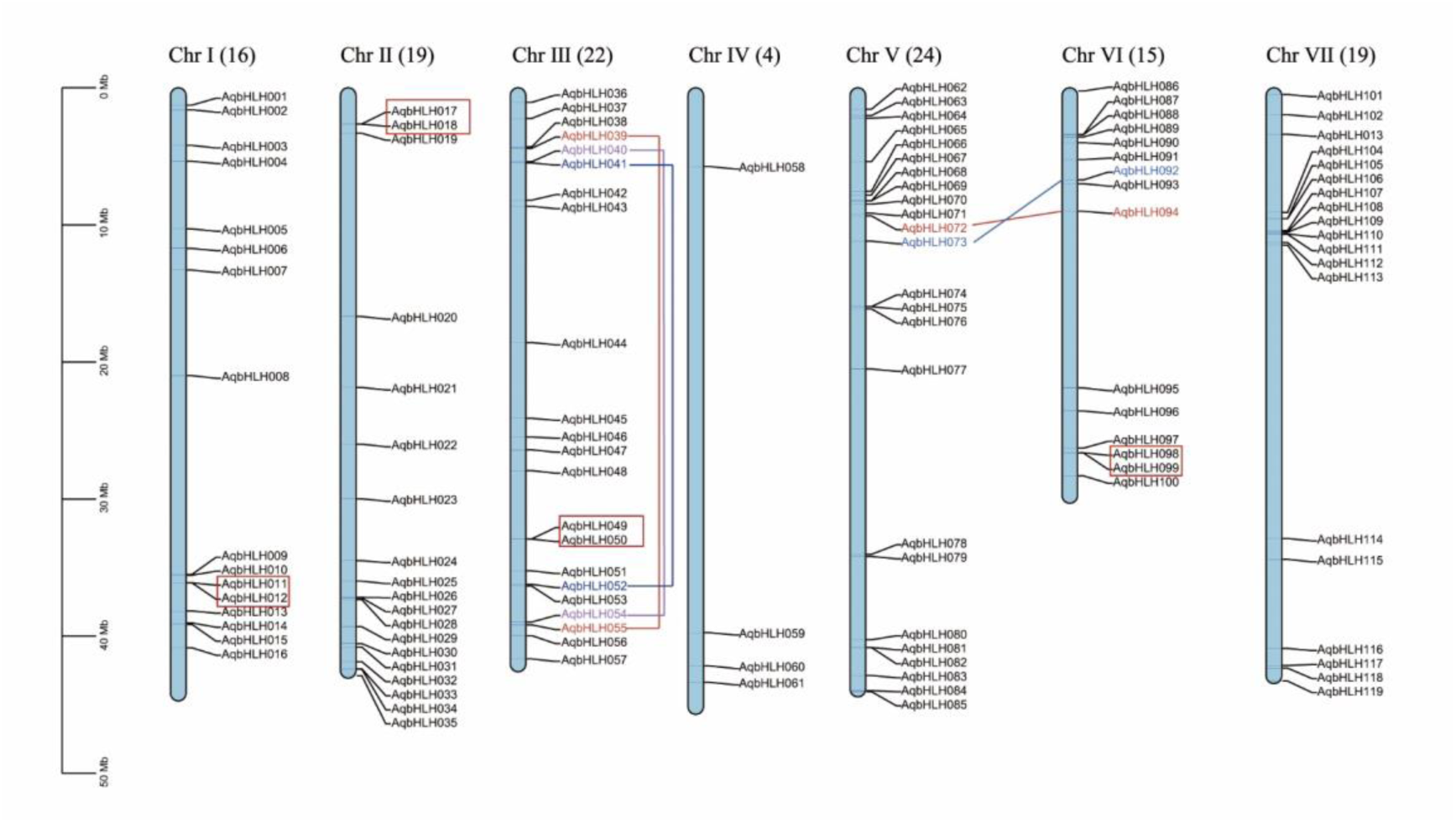
Chromosomal locations and duplication events for *AqbHLH* genes. The scale is in megabases (Mb). The total number of bHLH genes per chromosome is showed at the top of each chromosome. Segmentally duplicated genes in the chromosomes were indicated by color lines. Tandem duplicated genes were represented by red border.

Analysis of the genomic tandem arrays and intrachromosomal and interchromosomal duplicated regions, 18 genes of 120 *AqbHLH* genes showed syntenic relationships, occurring on chromosomes Ⅰ, Ⅱ, Ⅲ, Ⅴ and Ⅵ. The chromosome Ⅲ had eight genes showing syntenic relationships and belonging to S11 and S15 (**Figure 2 and 5**). Moreover, eight genes from four syntenic pairs underwent tandem duplications, including *AqbHLH 011*, *012*, *017*, *018*, *049*, *050*, *098* and *099*. *AqbHLH011* and *AqbHLH012* are the members of S8, and the *AqbHLH049* and *AqbHLH050* belong to S7, both of which comprised the most expended *AqbHLH* genes (**Figure 1**). And ten genes can be grouped into putative intrachromosomal and interchromosomal duplication events (**Figure 5**). Genes with syntenic relationships have similar protein structures and intron insertions (**Figure 3**).

### 3.5 Expression profiles of *AqbHLH* genes in spur and blade

In order to explore the functions of *AqbHLH* genes in the formation and development of spur in *A. coerulea*, we compared the *AqbHLH* genes expression levels in the spur cup (tubular structure at bottom of petal, **Figure 7C**) and blade (lamellar structure at top of petal, **Figure 7C**) at the 1, 3 and 7 mm developmental stages by using public RNA-Seq data [22]. Fifty-seven of all 120 *AqbHLH* genes were considered to be expressed with TPM value > 1 and were used in the following analysis (**Supplementary Table 4**). The expression patterns of *AqbHLH* genes were clustered into six main blocks. The *AqbHLHs* in block Ⅰand Ⅲ showed tissue-specific expression patterns, and constitutive expression with high expression levels during development in blade and spur, respectively. The genes in block Ⅳ and Ⅴ had higher expression levels in blade at 1 mm and spur at 7 mm (**Figure 6A**). We identified eight *AqbHLH* genes were differentially expressed genes (DEGs) between spur and blade at 1 and 3 mm developmental stages. The *AqbHLH028*, *AqbHLH046* and *AqbHLH082* were higher expressed in spur, contrary to *AqbHLH019, AqbHLH027*, *AqbHLH083*, *AqbHLH092* and *AqbHLH102* which showed higher expression levels in blade (**Figure 6A, Supplementary Figure 1A**). The different expression patterns of these DEGs might have close relationship with the special morphological characters of spur and blade.

**Figure 6.**
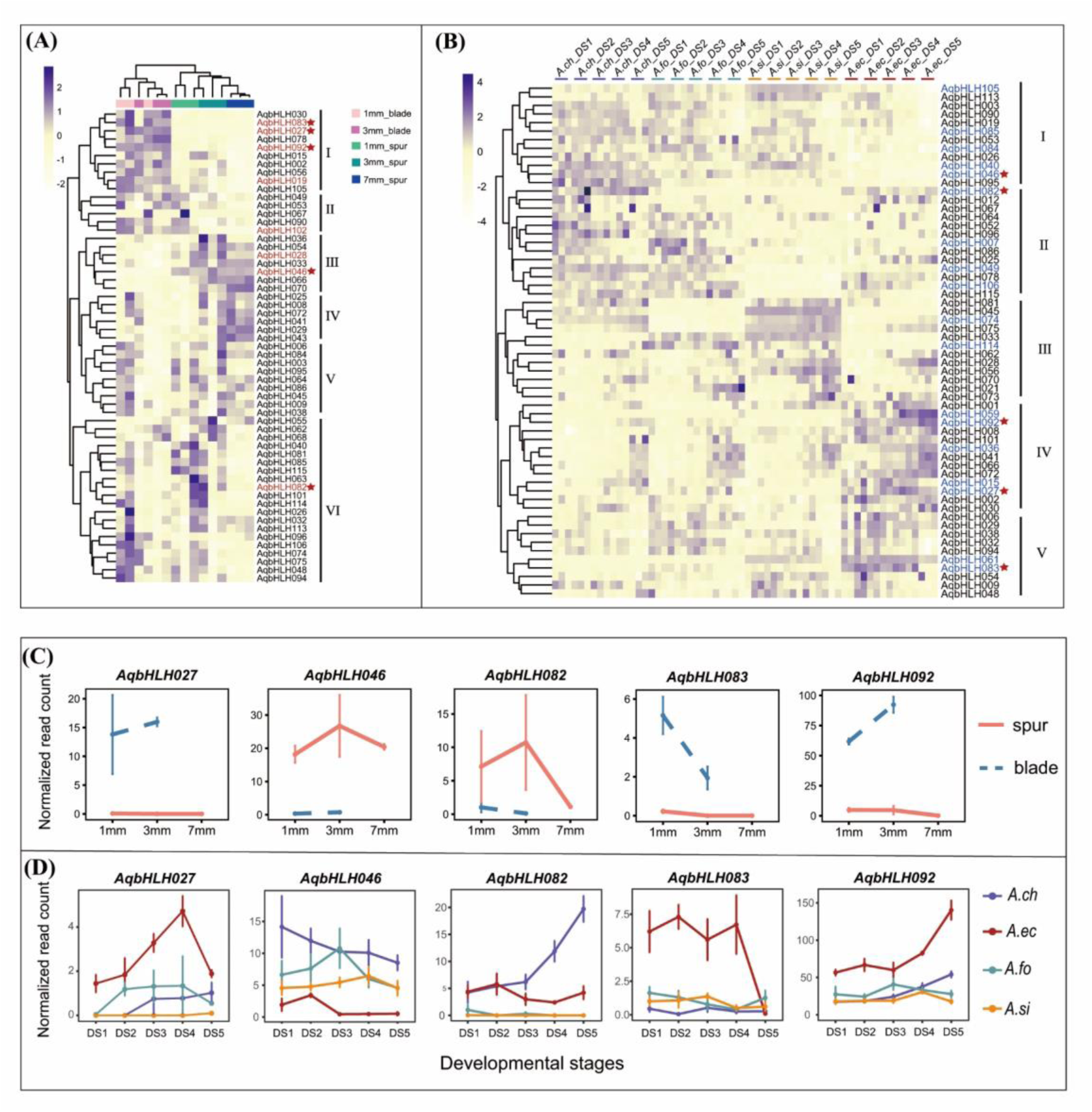
The expression of *AqbHLHs*. **(A)**, the expression pattern of *AqbHLHs in A.coerulea*. Red texts indicate the DEGs between spur and blade at 1mm and 3mm stages. **(B)**, the expression pattern of *AqbHLHs* in four species of *Aquilegia*. Blue texts indicate the common DEGs between *A.ecalcarata* (spurless taxa) and other species (spurred taxa) at DS1-DS5. The red five-pointed stars indicate the common DGEs in (A) and (B). **(C)** and **(D)**, the expression patterns of five common DEGs. *A.ch*=*A.chrysantha, A.fo=A.formosa, A.si=A.sibirica*. The gene expression level was normalized by using TPM.

**Figure 7.**
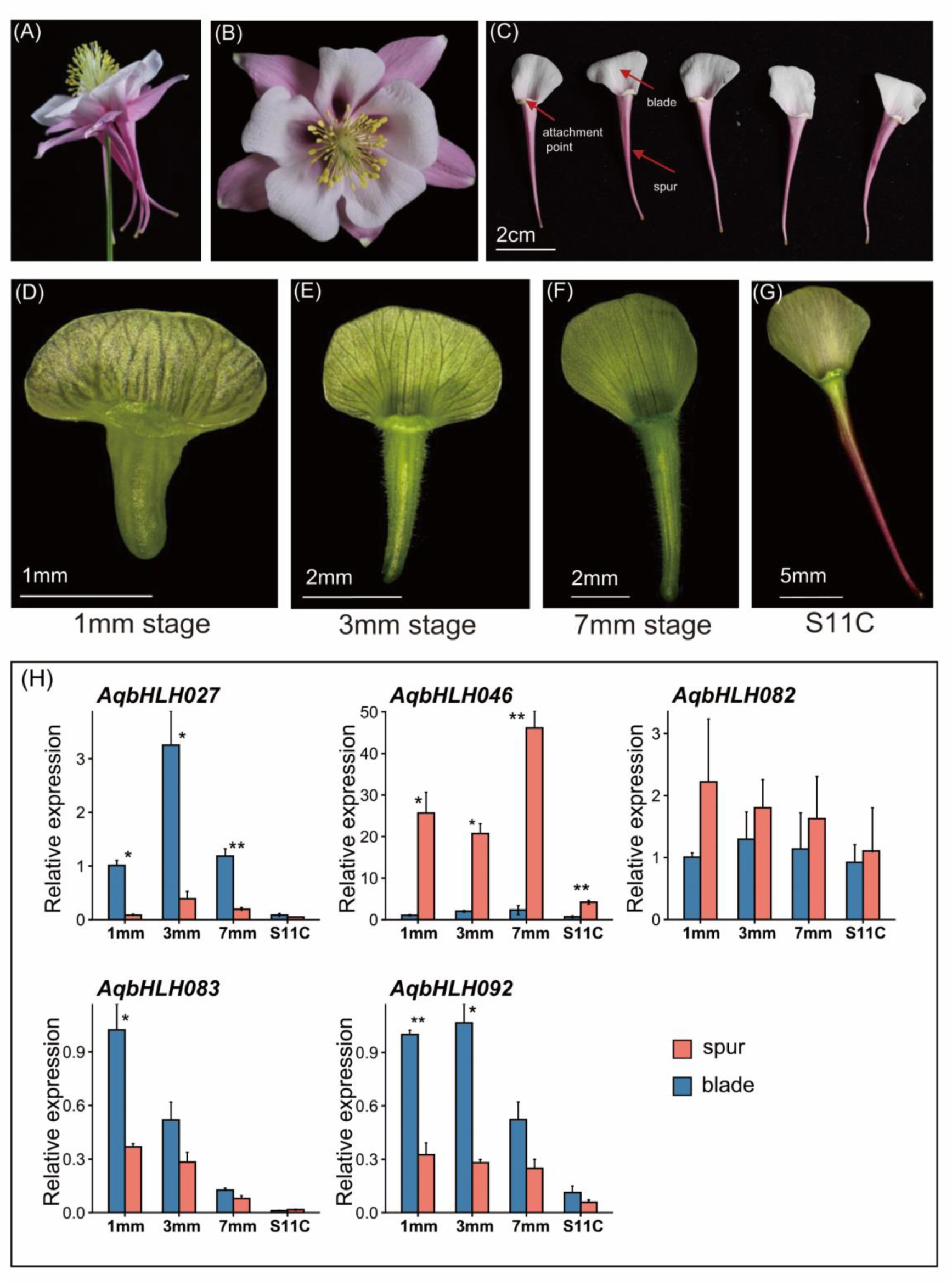
The petal structure of *A.coerulea* and qRT-PCR of five *AqbHLHs*. **(A)-(C),** the petal structure of *A. coerulea*. (**D)-(G)**, the morphology of petal in 1mm, 3mm, 7mm and S11C stage. (**H)**, the qRT-PCR patterns of five *AqbHLHs* in different developmental blade and spur at different developmental stages.

### 3.6 Expression profiles of *AqbHLH* genes in spurless and spurred species

To explore the relationship of *AqbHLH* genes expression and the formation and development of spur in *Aquilegia*, we compared these genes expression patterns in spurless taxa (*A. ecalcarata*) and spurred taxa (*A. chrysantha, A. formosa, A. sibirica*) at the early developmental stages of petals, which are indiscernible pre-spur formation at developmental stage 1 (DS1) and differentiated at stage 5 (DS5) by using public RNA-Seq data. The details of DS1-DS5 were described in Ballerini (Ballerini et al. 2019). Sixty *AqbHLH* genes were considered to be expressed with TPM value > 1 (**Supplementary Table 5**). The expression patterns of *AqbHLHs* were clustered into five main blocks. We found that these genes in blockⅠwere higher expressed in three spurred species than that in spurless taxa (*A. ecalcarata*), especially at DS4 and DS5. On the contrary, these genes in blocks Ⅳ and Ⅴ were higher expressed in spurless taxa. These genes in block Ⅱ and Ⅲ had no specific expression patterns between spurless and spurred taxa (**Figure 6B**).

Eighteen *AqbHLH* genes were identified as DEGs between spurless and spurred taxa at DS1-DS5 stages. The *AqbHLH015, AqbHLH027*, *AqbHLH059, AqbHLH061, AqbHLH083* and *AqbHLH092* were higher expressed in spurless taxa. While *AqbHLH040, AqbHLH046, AqbHLH084, AqbHLH085, AqbHLH105* and *AqbHLH114* were higher expressed in spurred taxa. (**Figure 6B, Supplementary Figure 1B**). The other six DEGs had no similar expression pattern in spurred taxa. For example, *AqbHLH082* was highest expressed in *A. chrysantha* followed by *A. ecalcarata*, and lowly expressed in *A. formosa* and *A. sibirica* at any stages. Notably, the expression patterns of the five differentially expressed *AqbHLHs,* including *AqbHLH027*, *AqbHLH046, AqbHLH082, AqbHLH083* and *AqbHLH092,* shared between different tissues in *A. coerulea* and between spurless and spurred species, as showed in **Figure 6C and D**. Significantly, the *AqbHLH046* was not only highly expressed in spur, but also showed higher expression levels in spurred species than that of spurless specie, which might suggest its important role in the regulation of formation and development of spur. The expression pattern of *AqbHLH092* is just opposite to that of the *AqbHLH046*, showing significantly higher expression level in blade and spurless species.

### 3.7 Expression validation of *AqbHLH* genes using qRT-PCR

To verify the expression patterns of *AqbHLHs*, the five differentially expressed *AqbHLHs* shared between different tissues and between different species were analyzed using qRT-PCR. We collected spur and blade at phase I (1, 3, 7 mm stages) and Phase II (S11C) of *A. coerulea* (**Figure 7A-G**). Consistent with RNA-Seq data, the *AqbHLH027, AqbHLH083* and *AqbHLH092* were significantly highly expressed in blade than in spur at four stages, except for *AqbHLH083* at S11C (**Figure 7H**). It is notable that the *AqbHLH046* was significantly higher expressed in spur than blade at four stages, especially at early stages 1 ∼ 3 mm, in line with the result of RNA-Seq analysis. *AqbHLH082* was also highly expressed in spur than in blade. Additionally, other 15 *AqbHLHs* belonging to blocks Ⅰ, Ⅲ and Ⅳ in figure 6 were selected and analyzed using qRT-PCR. The expression of *AqbHLH028* was higher in spur than in blade at four stages, in line with the result of RNA-Seq analysis (**Supplementary Figure 2**). Similarly, the expression of *AqbHLH036* and *AqbHLH066* were highly expressed in spur than in blade at four stages, especially at S11C. The results showed that there was a faint difference in the expression of *AqbHLH002*, *AqbHLH015*, *AcoebHLH019*, *AqbHLH029*, *AqbHLH030*, *AqbHLH054*, *AqbHLH056*, *AqbHLH070 AqbHLH102* and *AqbHLH105* between blade and spur at any stage, which was consistent with the result of RNA-Seq analysis (**Supplementary Figure 2**).

### 3.8 Protein-protein interactions (PPI) and co-expression analysis for *AqbHLHs*

To probe the interactive relationship of the *AqbHLH046* and *AqbHLH092,* two potential key regulators, with other TFs and proteins, we constructed networks of PPI network map by using the homology in Arabidopsis. The results of this prediction indicated that *AqbHLH046* (*AT5G50915/bHLH137* ortholog) interacted with four bHLH TFs, including *AqbHLH027* (*MUTE/bHLH45* ortholog), *AqbHLH053* (*PYE/bHLH47* ortholog), *AqbHLH060* (*AT3G20640/bHLH123* ortholog) and *AqbHLH083* (*MUTE/bHLH45* ortholog) and four MYB TFs, including *AqMYB60* (*MYB60* ortholog), *AqMYB105_1, AqMYB105_2* and *AqMYB105_3* (*MYB105* ortholog), *AqbHLH092* (*bHLH63/CIB1* ortholog) interacted with *AqCRY2* (*CRY2* ortholog) (**Figure 8A**). Furthermore, we analyzed the co-expression for these genes in *A. coerulea*. The expression level of *AqbHLH046* was a significantly positive correlation with *AqMYB105_3* and negative correlation with *AqbHLH027, 083, 092* and *AqCRY2*. The expression level of *AqbHLH092* was positive correlation with *AqCRY2*, *AqbHLH027* and *AqbHLH083* and significantly negative correlation with *AqMYB105_1*, *105_2*, *105_3* and *AqbHLH046* (**Figure 8B**). Furthermore, we found the expression of *AqbHLH046* was significantly positively (R = 0.75, p < 0.001) correlated with *POPOVICH* which has been proved to play a central role in the development of spur by regulating cell division in *A. coerulea* (**Figure 8C**).

**Figure 8.**
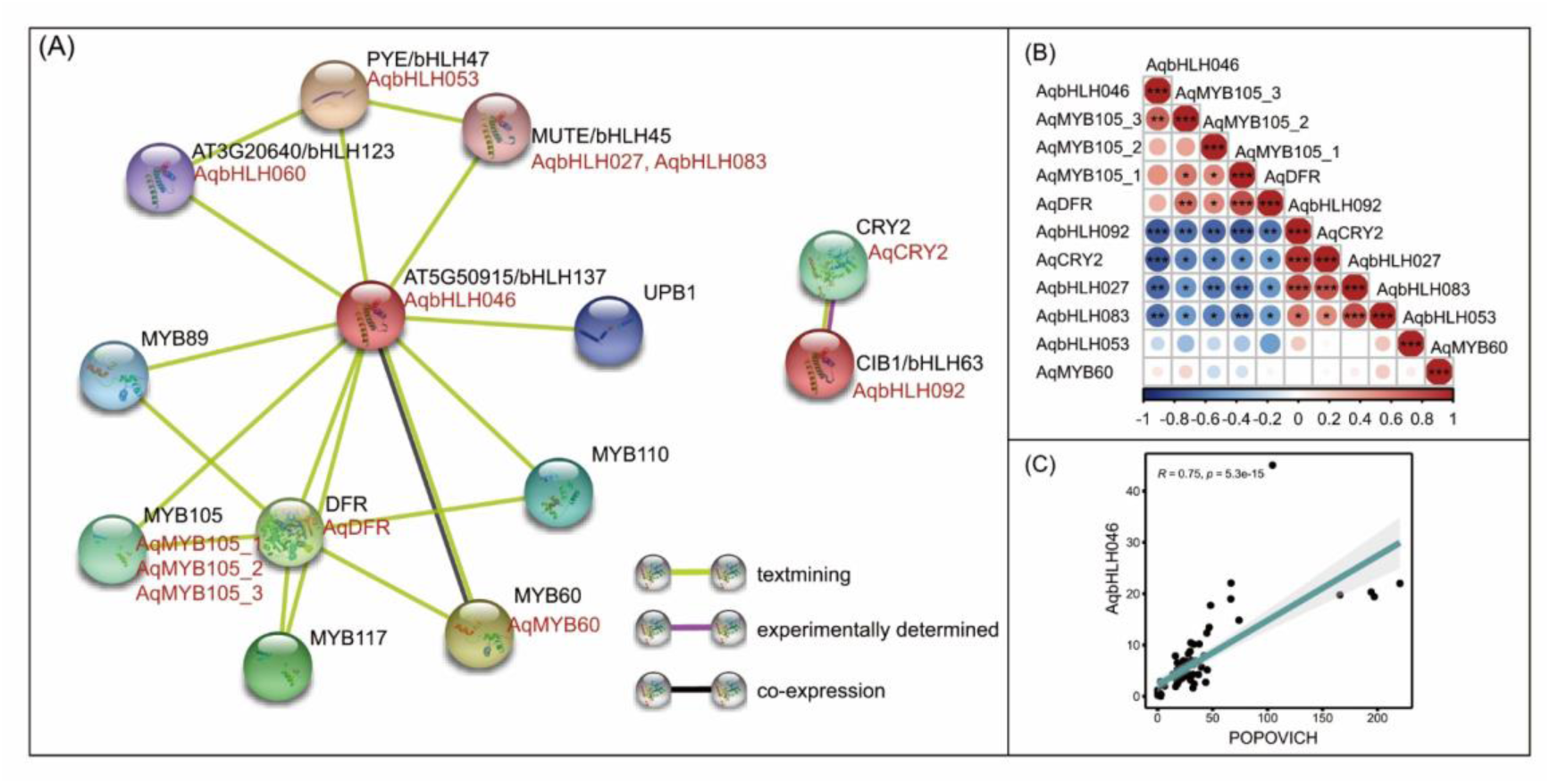
(A), protein-protein interaction network for key *AqbHLHs* based on their orthologs in *Arabidopsis*. The black texts and red texts represent the proteins in *Arabidopsis* and *AqbHLH* proteins, respectively. **(B),** the coexpression of genes in (A). **(C),** the coexpression of *POPOVICH* and *AqbHLH046*.

## 4. Discussion

### 4.1 The bHLH TFs family in genome of *A. coerulea*

The *bHLH* TFs family is one of the largest families of TFs in plants and was involved in plant growth, development, and stress responses (Ito et al., 2012; Li et al., 2019; Arai et al., 2019; Kavas et al., 2016). Thus they were well studied in many plants, especially in model plants or agricultural crops. For example, two atypical bHLH proteins *POSITIVE REGULATOR OF GRAIN LENGTH 1 (PGL1)* and *POSITIVE REGULATOR OF GRAIN LENGTH 2 (PGL2)* and typical bHLH protein *ANTAGONIST OF PGL1 (APG)* are involved in controlling rice grain length and weight (Heang and Sassa, 2012a, 2012b). *Aquilegia* species are famous horticultural plants with abundant color and floral morphology. It is essential to reveal the regulation mechanism of floral morphology, such as the development of petal nectar spur, for new variety breeding and adaptive evolution research. However, the *bHLH* TFs in *A. coerulea* have not been reported. In this study, we identified 120 *bHLHs* with complete domains in *A. coerulea,* which is less than that in *Helianthus annuus* (183), *A. thaliana* (162), and rice (167) (Heim et al., 2003; Sun et al., 2015; Li et al., 2021b), and more than that in sweet cherry (66) and *Nicotiana tabacum* (100) (Bano and Patel, 2021; Shen et al., 2021). Given that the genome size of *A. coerulea* (∼300M) is larger than *A. thaliana* (∼125M) and smaller than rice (∼466M) and *N. tabacum* (∼4.5G), the difference in the number of identified *bHLH* genes among these species is related to genome size as suggested by Li et al (2021). We speculated that the difference between the number of identified *bHLH* genes among these species is at least partly attributed to the quality and complexity of genome sequence. The size of the assembled genome accounts for only 77.8% of the estimated sweet cherry genome size (Shirasawa et al., 2017). The assembled genome of *A. coerulea* used in this study is smaller than previous estimates (∼ 300Mb versus ∼ 500Mb) (Bennett et al., 1982; Bennett and Leitch, 2011), is likely due to gaps and repetitive content that significantly affects the assembly quality (Filiault et al., 2018). Additionally, gene duplications are considered to be one of the primary driving forces in the expansion and evolution of a gene family, in turn, often indicates new gene functionalization (Ma et al., 2008; Ke et al., 2020). In *A. thaliana* and rice genome, 48% (73/147) of the *AtbHLH* proteins and 48% (80/167) of the *OsbHLH* proteins might have evolved from segmental duplications, tandem duplications, and polyploidy duplications (Toledo-Ortiz et al., 2003; Bailey et al., 2003; Li et al., 2006). These genes duplication might result in abundant bHLH proteins in *A. thaliana* and rice genome. In our study, there are 120 identified *AqbHLH* proteins in *A. coerulea,* and 18 of all *AqbHLH* (15%) genes might have evolved from genome duplicated events. The fewer genes duplication could be one of factors leading to fewer *AqbHLH* genes in *A. coerulea*.

Sequence analysis revealed that each putative *AqbHLH* protein contains four conserved regions including a basic region, a loop region and two helixes (**Figure 1**). The four conserved regions were highly conserved in the majority of the *AqbHLH* genes, as well as in *A. thaliana*, sunflower and spine grapes (Bano and Patel, 2021; Li et al., 2021b). The bHLH domain is important for DNA binding and dimerization (Murre et al., 1989), which contained Leu-23, Pro-28, Tyr-47, Leu-51 and Leu-58 with conservation of more than 90% in *A. coerulea*. Moreover, Leu-27 in helix I (correspond to Leu-23 in *AqbHLH* genes) and Leu-73 in helix II (correspond to Leu-51 in *AqbHLH* genes) were crucial for the interaction of proteins and had a close relationship with the function of *bHLH* genes (Carretero-Paulet et al., 2010), indicating their essential roles in bHLH proteins.

In plants, the *bHLH* family genes were divided into 15 ∼ 31 subfamilies (Wang et al., 2018a; Bailey et al., 2003; Ke et al., 2020). According to phylogenetic analysis, we classified 120 identified *AqbHLH* genes into 15 subfamilies (**Figure 2**) and found that the members of *AqbHLH* proteins in S7 and S8 subfamilies occurred marked expansion. The homologous gene in *A. thaliana* (*At1G49770.1/ RGE1*) in S7 subfamily has been reported to control embryo growth from the endosperm that plays an important role in driving rapid barriers between divergent species by facilitating incompatible interactions (Kondou et al., 2008; Kinser et al., 2021). The *At1G10610.1* (*AtbHLH95*) in S8 subfamily regulates embryo growth, endosperm breakdown and embryonic epidermal development (Kondou et al., 2008; Yang et al., 2008). We speculated that the homologous genes of *AtRGE1* and *AtbHLH95* in *AqbHLH*s may be contributed to the post-zygotic isolation by regulating the embryo growth that may be associated with maintaining independent among species in *Aquilegia*. However, that view remains to be proven.

Protein motif and gene structure analysis showed that the members of the same subfamily generally contain similar protein motifs and show conserved gene structures. The types and composition of inner motifs mainly determine the protein function. Some motifs only could be detected in specific subfamilies, such as the motifs 3 and 4 in S7 subfamily, and the motifs 5, 10 and 15 in S8 subfamily, which might be related to their specific function of members in the two subfamilies. Compared with the conservative protein motif pattern, *AqbHLH* family has diverse intron/exon organizations in the same subfamily and among different subfamilies. The phenomenon also was found in the *bHLH* gene family in Chinese white pear (Dong et al., 2021), sorghum (Fan et al., 2021) and tobacco (Bano and Patal, 2021). The result suggested that the *AqbHLH* gene family may play diverse roles in many biological processes. The diverse *cis*-acting elements in the promoter regions of *AqbHLH* genes also provide important cues for their diverse biological functions.

### 4.2 Petal spur development and bHLH genes

The functions of *bHLH* transcription factors have been reported in animals and plants, including control of cell proliferation and specific cell lineages development (Heim et al., 2003; Toledo-Ortiz et al., 2003; Pillitteri et al., 2007; Pires and Dolan, 2010; Girin et al., 2011; Zhang et al., 2018; Zhang et al., 2021; Nan et al., 2022). Spur as a specific adjacent tissue, the pattern of cell division and cell lineages are different from the blade in *Aquilegia* (Shan et al., 2019; Yant et al., 2015). According to the morphological studies of spur development in *Aquilegia*, when spur length is less than 10 mm, cell division is a major contributor to the early development of spur growth (Puzey et al., 2012; Yant et al., 2015). We identified five *AqbHLH* genes, namely *AqbHLH027*, *AqbHLH046*, *AqbHLH083* and *AqbHLH092*, were differentially expressed between spur and blade at early developmental stages (1 and 3 mm stages) in *A*. *coerulea*. Furthermore, they were also differentially expressed between flowers of spurred and spurless *Aquilegia* species at early developmental stages (DS1-DS5). Based on expression patterns and functions of homology, we inferred that they might be involved in spur formation by regulating related to different cell activities including meristem cell division, cell differentiation and cell fate in phase I of spur development. First, the *AqbHLH027* and *AqbHLH083* shared a high degree of homology with the *MUTE*, which disturbs the asymmetrically stomatal cell division and differentiation. In the absence of *MUTE*, the cells occur excessive divisions and fail to differentiate stomata in *Arabidopsis* (Pillitteri et al., 2007). Nectar spur development is related to the re-initiation of the meristematic activity program which is similar to stomata differentiation. The expression levels of *AqbHLH027* and *AqbHLH083* were lower in spur than that of blade in *A. coerulea* (**Figure 6C**), and were also lower in spurred species (**Figure 6D**). The cell division is concentrated in spur which might attribute to the lower expression levels of *AqbHLH027* and *AqbHLH083*.

Second, the *AqbHLH046* showed a high degree of homology with *CKG* (*CYTOKININ-RESPONSIVE GROWTH REGULATOR*), which is involved in CK-dependent regulation of cell expansion and cell cycle progression by regulating the expression of *WEE1* (Park et al., 2021). The gene was significantly higher expressed in spur and spurred taxa in our study (**Figure 6C-D; Figure 7H**). Moreover, the protein-protein interaction analysis indicated that the expression level of *AqbHLH046* was significantly positive correlative with *POPOVICH* which has been proved to play a central role in the development of spur by regulating cell division in *A. coerulea,* and negative correlation with *AqbHLH027* and *AqbHLH083* (**Figure 8B**). The *AqbHLH046* also interacted with several *MYB* TFs (*AqMYB60, AqMYB105_1, AqMYB105_2, AqMYB105_3*) (**Figure 8A-B**). The *MYB105* was involved in boundary specification, meristem initiation and maintenance, and organ patterning (Lee et al., 2009). So we speculated that the *AqbHLH046* plays a key role in regulating organ size and spur development in *A. coerulea* at an early stage by regulating the cell expansion and cell cycle progression.

Third, the *AqbHLH092* is homologous with *CIB1* and exhibited a lower expression level in spur at 1-3mm stages (**Figure 6C; Figure 7H**) and spurless taxa at DS1-DS5 (**Figure 6D**). *CIB1* often interacts with *CRY2* to regulate growth, floral initiation and development. Such as, *CRY2* interacts with *CIB1* to regulate floral initiation and flowering in *Arabidopsis*, and *OsCIB1*-*OsCRYs* controls leaf sheath length via the GA-responsiveness (Liu et al., 2008; Li et al., 2021a). Additionally, *CIB1-CRY2* inhibits proteins that modulate cytoskeleton and cell cycle living mammalian cells (Park et al., 2016). In our study, we found the expression of *AqCIB1* and *AqCRY2* were positively correlated and were lowly expressed in spur (**Figure 8B**). We speculated that less *AqCIB1* and *AqCRY2* would likely involve spur development by controlling the cell cycle.

In addition to cell division, *AqbHLH036* was highly expressed at Phase II of spur development (S11C stage) in *A. coerulea* where cell mitotic activity ceases and the elongation of the differentiating cell (**Supplementary Figure 2**). *AqbHLH036* (ortholog of *RHD6*) was specifically involved in root hair cells elongation (Feng et al., 2017). Which suggested that the gene might be related to cells elongation. In spurless and spurred *Aquilegia* species, the expression levels of *AqbHLH036* showed an increasing trend with the development of flower in four *Aquilegia* species and were highly expressed in spurless taxa than in spurred taxa (**Supplementary Figure 1**). That might be implied the transition from cell division to cell expansion of spurless taxa is earlier than in spurred taxa, which contributed to the difference of spur length between them (Ballerini et al., 2019). Genome-wide identification and expression analysis of bHLH transcription factors in *Aquilegia* provide a theoretical foundation and valuable insights for spur formation and development.

## 5. Conclusion

*Aquilegia* species are an ideal flora model system for the evolution because of the remarkable diversity of shape in petal nectar spur. Here, 120 *AqbHLH* genes in the *A. coerulea* ‘Goldsmith’ genome were identified and divided into 15 subfamilies. The members of subfamily S7 were expansive, and they were related to the development of endosperm in the embryo. Notably, while *AqbHLH027, AqbHLH083* and *AqbHLH092* were higher expressed in blade, *AqbHLH046* was highly expressed in spur. They would be associated with spur development by regulating cell division and cell cycle as the primary or minor factors in phase I. Otherwise, *AqbHLH036* was likely to be involved in the cell elongation in spur growth during phase Ⅱ by controlling the transition from spur cell division to spur cell expansion. Our results provide the potential for discovering new genes that give rise to the petal nectar spur in *Aquilegia*.

## AUTHOR CONTRIBUTIONS

Xueyan Li, Hui Huang, and Zhi-Qiang Zhang conceived and performed the original research project. Xueyan Li performed the experiments, analyzed the data, and wrote the original draft. Hui Huang and Zhi-Qiang Zhang supervised the experiments and revised the writing, and obtained the funding for the research project. All authors read and approved the final manuscript.

## ACKNOWLEDGMENTS

We thank State Key Laboratory for Conservation and Utilization of Bio-Resources in Yunnan and Yunnan Key Laboratory of Plant Reproductive Adaptation and Evolutionary Ecology, School of Ecology and Environmental Sciences, Yunnan University, Yunnan, Kunming, China for providing the experimental facility.

## Captions

**Supplementary Figure 1 | (A)**, The expression patterns of 8 DEGs between spur tissue and blade tissue at 1mm stage and 3mm stages. **(B)**, The expression patterns of common 18 DEGs between *A.ecalcarata* (spurless taxa) and other species (spurred taxa) at DS1-DS5.

**Supplementary Figure 2 |** The qRT-PCR patterns of fifteen *AqbHLHs* in the different developmental blade and spur tissues at different developmental stages.

**Supplementary Table 1 |** Information of elements for *AqbHLHs*.

**Supplementary Table 2 |** Primers of qRT-PCR for *AqbHLHs*.

**Supplementary Table 3 |** Features of *AqbHLH* genes identified in *A.coerulea*.

**Supplementary Table 4 |** The TPM matrix of *AqbHLHs* in *A.coerulea*.

**Supplementary Table 5 |** The TPM matrix of *AqbHLHs* in four species of *Aquilegia*.

