## Supplementary figures and images for "Genome-wide identification and expression analysis of bHLH transcription factors reveal their putative regulatory effects on petal nectar spur development in *Aquilegia*"

### Fig S1

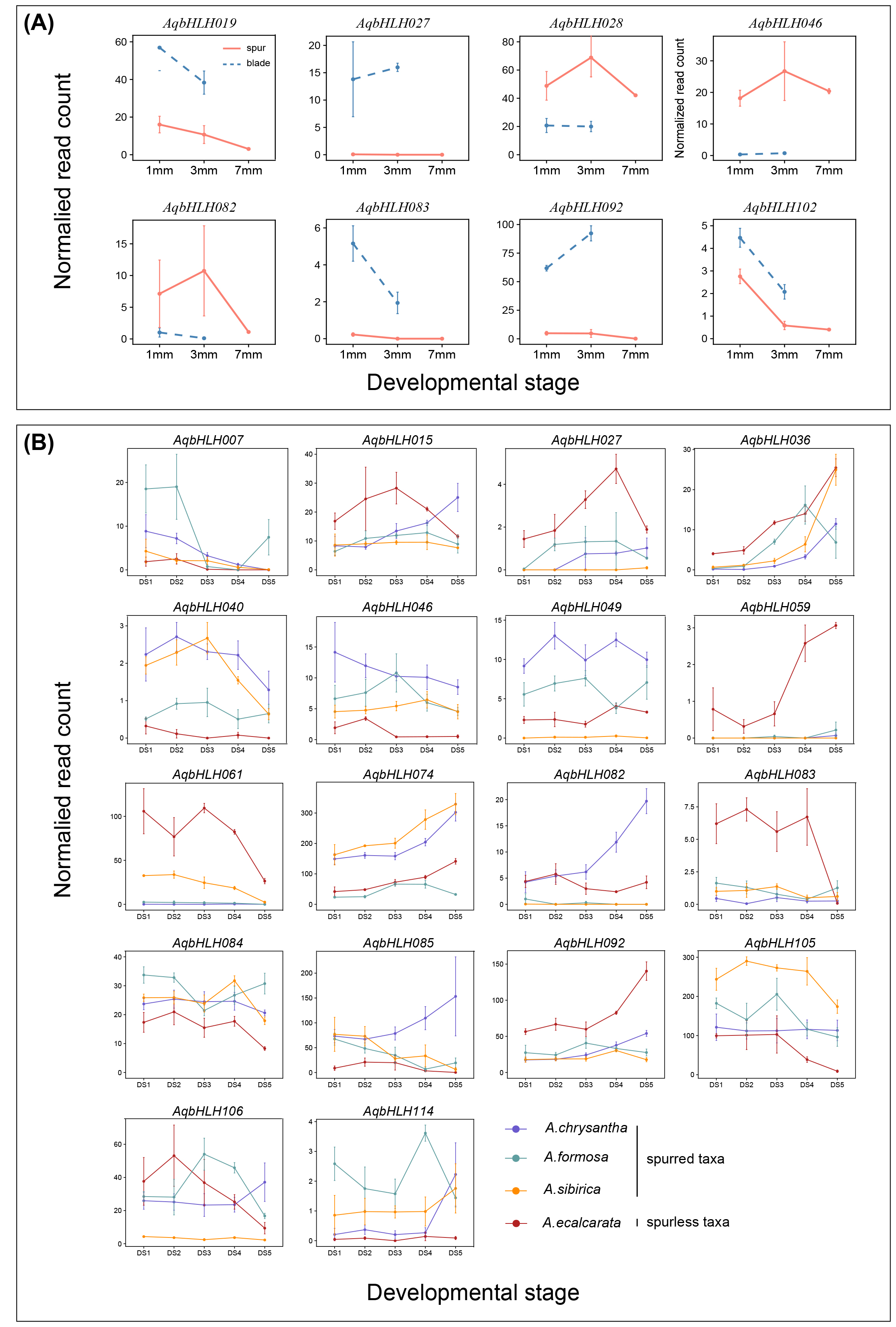

### Fig S2

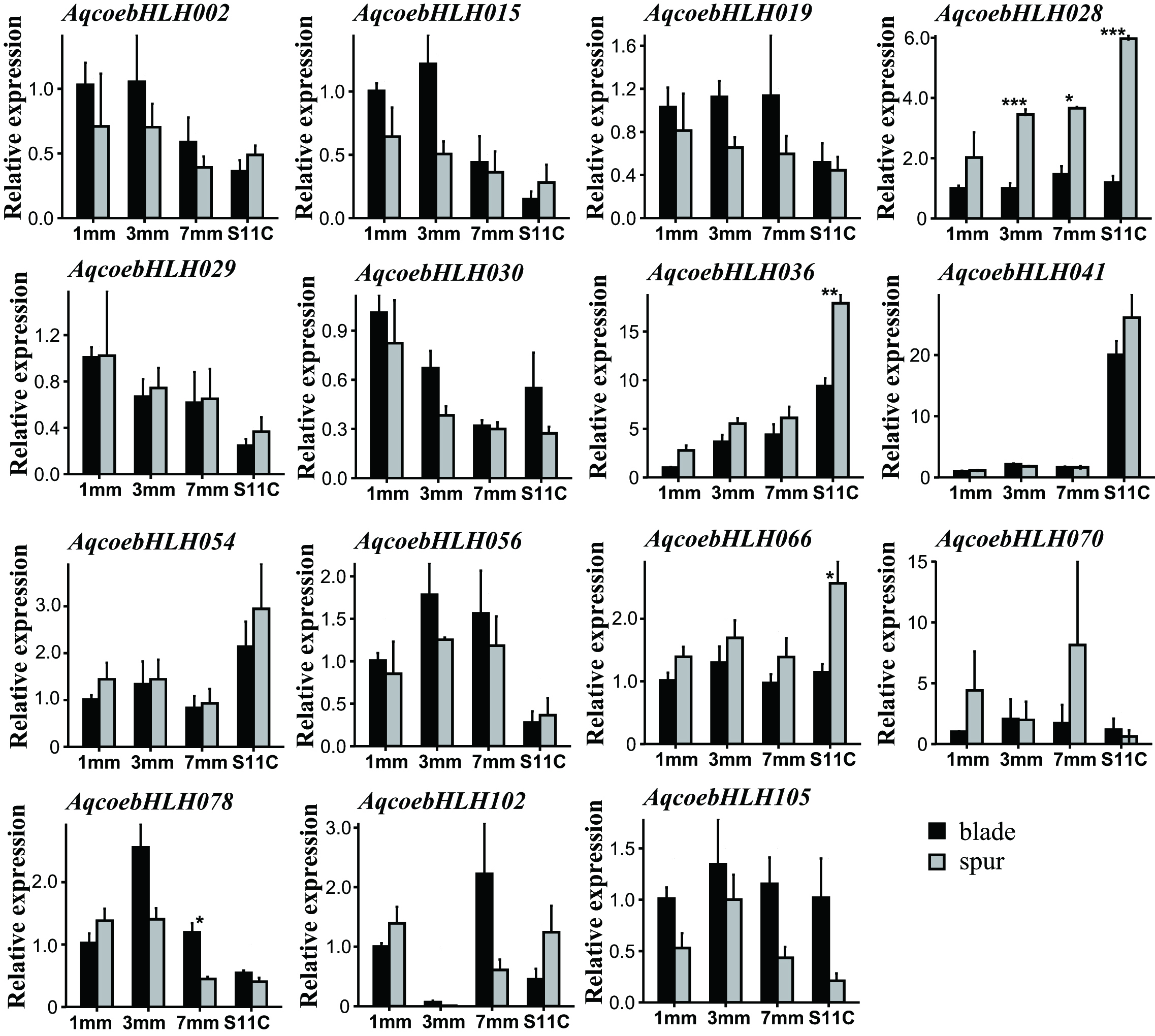
